# MALVIRUS: an integrated web application for viral variant calling

**DOI:** 10.1101/2020.05.05.076992

**Authors:** Simone Ciccolella, Luca Denti, Paola Bonizzoni, Gianluca Della Vedova, Yuri Pirola, Marco Previtali

## Abstract

Being able to efficiently call variants from the increasing amount of sequencing data daily produced from multiple viral strains is of the utmost importance, as demonstrated during the COVID-19 pandemic, in order to track the spread of the viral strains across the globe.

We present MALVIRUS, an easy-to-install and easy-to-use web application that assists users in two tasks:

1. computing a variant catalog consisting in a set of population SNP loci from the population sequences and
2. efficiently calling variants of the catalog from a read sample.

Tests on Illumina and Nanopore samples prove the efficiency and the effectiveness of MALVIRUS in genotyping SARS-CoV-2 strain samples with respect to GISAID data.

## Introduction

The SARS-CoV-2 pandemic has put the global health care services to the test and many researchers are racing to face its swift and rapid spread. Since the outbreak of the virus in China and in other European countries, several studies are using sequencing technologies to track the geographical origin of SARS-Cov-2 and to analyze the evolution of sequence variants (1, 2). In this context, the availability of efficient approaches to analyze variations from the growing amount of sequencing data daily produced is of the utmost importance.

The typical pipelines for the analysis of variations within viral samples consists in aligning reads against a reference genome (3), then analyzing the alignments to discover the variants (4, 5). However, the increasing number of viral assemblies available in public databases such as GISAID (6), GenBank (7), and the COVID-19 Data Portal allows to build a complete catalog of variants of a viral population. Such a catalog can be used to reduce the complexity of comparative analysis of genetic variants of sequencing samples. Clearly, to this aim, it is crucial that users are assisted by an efficient and easy-to-use method for building and updating the catalog and for calling variants that are in this catalog. In this paper, we introduce MALVIRUS, a web application for quickly genotype newly sequenced viral strains, including but not limited to the SARS-CoV-2 strains. The application is distributed as a multi-platform Docker container and it can be easily accessed using any modern Internet browser. As use case, we show that MALVIRUS is accurate at genotyping newly sequenced SARS-CoV-2 strains on both short and long read data.

## Methods

To efficiently genotype a viral sample from an individual with respect to the current knowledge, we propose MALVIRUS a web application based on five state-of-the-art tools.

The application is divided into two logically distinct modules: the creation of the catalog containing the SNP loci of the viral species under investigation and the variant calling from the read sample. Fig. 1 shows the MALVIRUS pipeline.

**Fig. 1.**
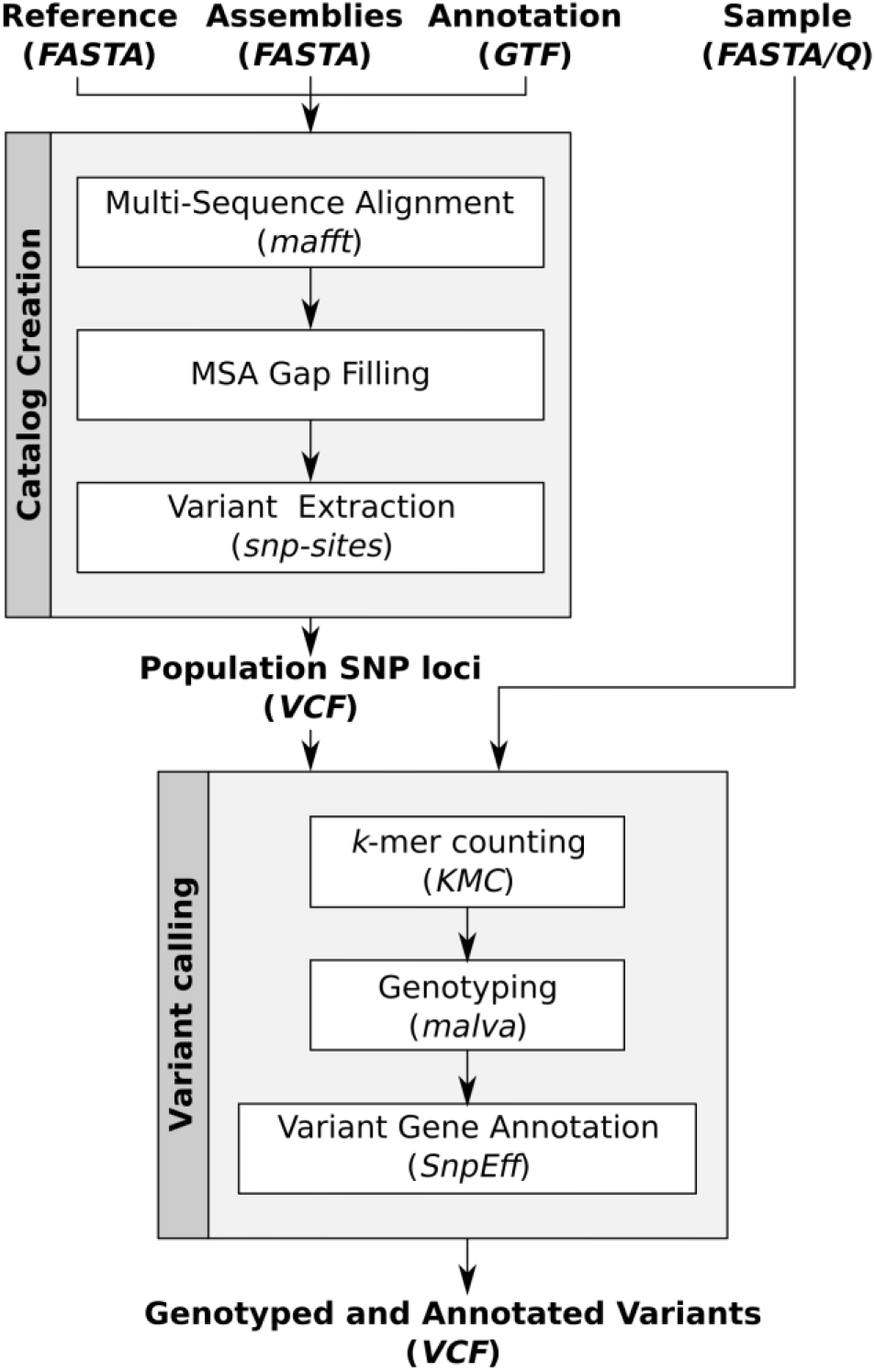
Schematic representation of the pipeline integrated in MALVIRUS.

The first module requires as input the reference genome of the species under investigation, the assemblies of a set of strains of that species, and, if available, the annotation of the genes. The output of this module is the set of population SNP loci in VCF format. MALVIRUS first builds the full-length sequence alignment of the input sequences to the input reference genome using MAFFT (8), then extracts the set of population SNP loci from the multiple alignment using snp-sites (9). Since snp-sites is not able to output variants in positions with gaps, MALVIRUS fills the gaps in the alignment with the corresponding portions of the reference. Although this step might induce some artificial variants, it allows to preserve real ones that might be lost due to incomplete assemblies. If the population under investigation is well characterized and/or the user wants a finer control over the variant catalog, it is possible to upload a custom catalog of SNP loci in VCF format instead of relying on the automatic computation from a set of assemblies.

The second module requires as input a sample of reads in FASTA/Q format and a catalog of population SNP loci along with the corresponding reference genome chosen among the ones computed or uploaded in the first module. The output of the second module is a VCF containing the genotype information of the sample and their qualities. To call the genotype of each variant, this module counts the *k*-mers in the sample using KMC3 (10), then it genotypes the variants using MALVA (11): an efficient and accurate mapping-free approach for genotyping a set of known SNPs and indels initially developed for human individuals. We extended MALVA to support haploid organisms and high-coverage samples. Additionally, if gene annotation is available, the module also annotates the functional effects of each variant using SnpEff (12). Finally, the results of each analysis can be visualized as a table (see Fig. 2 for an example) or downloaded in VCF format or as a spreadsheet for further analysis.

**Fig. 2.**
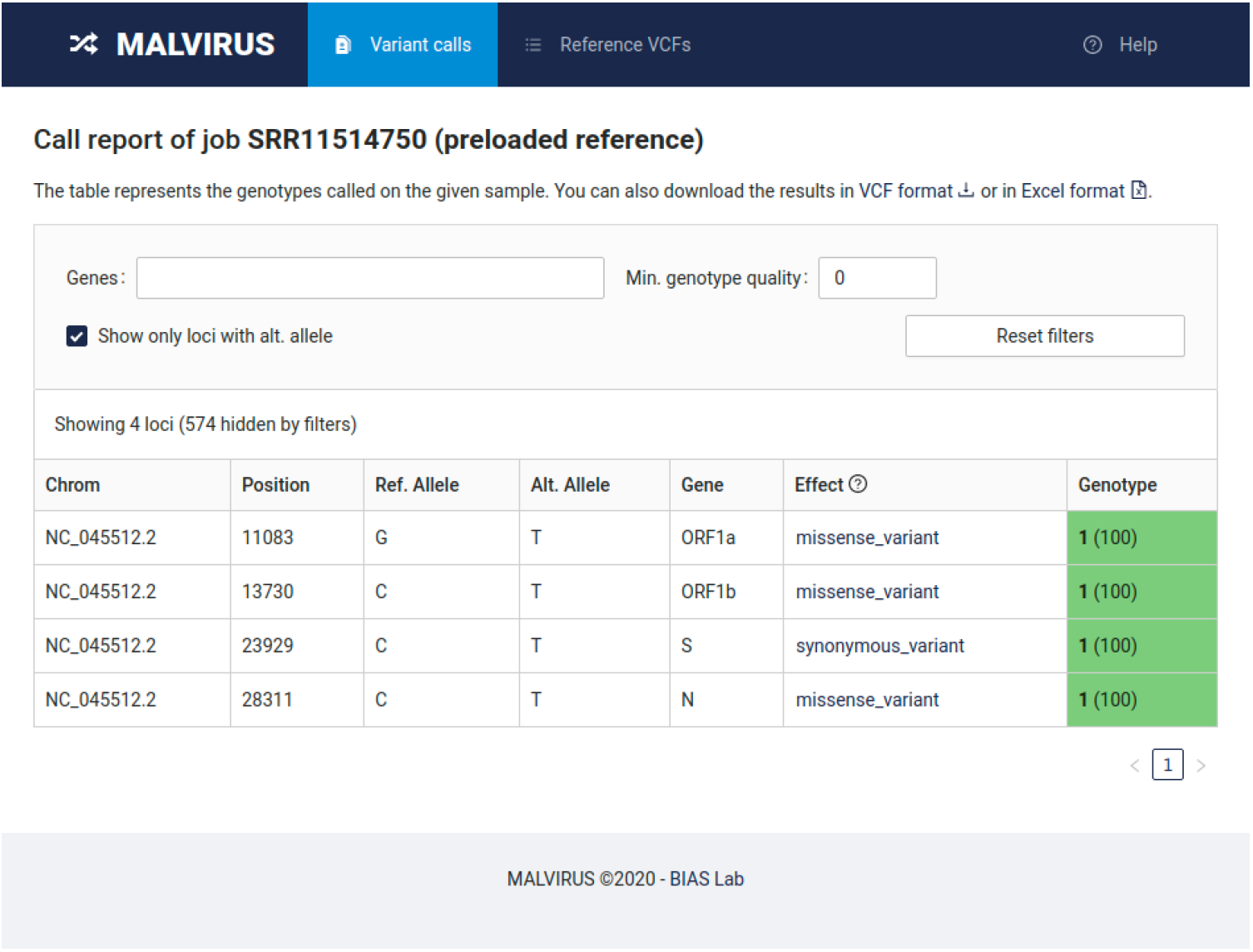
Example of the final report of MALVIRUS.

MALVIRUS is available as a self-hosted web application distributed as a Docker container image that can be installed and run on multiple platforms, from personal laptops to large cloud infrastructures. For user convenience, the application is distributed with a set of precomputed catalogs of variants for SARS-CoV-2 based on the assemblies available on GenBank (7), therefore the user can immediately run MALVIRUS on a locally available (*e.g*., private) viral sample. The precomputed catalogs can be easily updated from the application itself with a single click.

Extensive documentation and a detailed tutorial are available at https://algolab.github.io/MALVIRUS.

## Results

To test the effectiveness of MALVIRUS, we considered 10 strains from the GISAID database for which a sample of raw reads is available on the Sequence Read Archive (SRA). These were the only samples that we were able to cross-reference between GISAID and SRA at the moment of writing, furthermore for 5 of such strains we analyzed reads sequenced using both Illumina and Oxford Nanopore technologies, showing that MALVIRUS achieves similar results on both data types.

For simulating a real case scenario, where the goal is to genotype a newly-sequenced strain, before analyzing a sample, we removed it from the set of complete SARS-CoV-2 strains available on GISAID (accessed on July 17, 2020) and we ran MALVIRUS on the remaining 42709 strains for building the variant catalog. From the 42709 strains, the first module of MALVIRUS produced a VCF containing 13709/13710 variants (depending on which strains were removed). Then, we genotyped such a catalog using the second module of MALVIRUS starting from the corresponding read samples.

To evaluate the overall accuracy of MALVIRUS we computed its precision and recall in genotyping the set of known variants produced by its first module. To compute precision and recall, we used the first module of MALVIRUS to build the variant catalog with respect to the considered strain (*i.e.,* the strain we removed) and we used it as truth set. We then classified each variant as a *reference variant* if its real genotype is 0, *i.e.* the reference allele, and as an *alternate variant* if its genotype is not 0. Finally, we compute the precision and recall of MALVIRUS and reported the results of this analysis in Table 1.

**Table 1.** Results on real data. For each considered strain (GISAID ID, for ease of presentation we removed the *EPI_ISL_* prefix) and the corresponding SRA sample, we report the Precision and Recall obtained by MALVIRUS on calling reference variants (i.e. those variants whose real genotype is the reference allele, **REF**) and alternate variants (i.e. those variants whose real genotype is the alternate allele, **ALT**). For each sample, we also report the technology used (*ONT* for Oxford Nanopore and *ILLU* for Illumina) and its coverage (in terms of number of bases).

| *GISAID ID* | *SRA ID* | *Seq. Tech.* | *# of bases* | *Precision REF* | *Recall REF* | *Precision ALT* | *Recall ALT* |
| --- | --- | --- | --- | --- | --- | --- | --- |
| *416410* | SRR11397727 | ONT | 331 | 1 | 0.999 | 1 | 0.5 |
| *416410* | SRR11397730 | ILLU | 178 | 1 | 0.999 | 1 | 0.5 |
| *416411* | SRR11397726 | ONT | 363 | 1 | 1 | 1 | 1 |
| *416411* | SRR11397729 | ILLU | 109 | 1 | 1 | 1 | 1 |
| *416412* | SRR11397721 | ILLU | 126 | 1 | 1 | 1 | 1 |
| *416412* | SRR11397725 | ONT | 236 | 1 | 0.999 | 1 | 0.57 |
| *416413* | SRR11397720 | ILLU | 82 | 1 | 1 | 1 | 1 |
| *416413* | SRR11397724 | ONT | 221 | 1 | 0.999 | 1 | 0.83 |
| *416415* | SRR11397718 | ILLU | 112 | 1 | 0.999 | 1 | 0.6 |
| *416415* | SRR11397722 | ONT | 381 | 1 | 0.999 | 1 | 0.4 |
| *416514* | SRR11397717 | ILLU | 89 | 1 | 1 | 1 | 1 |
| *416515* | SRR11397716 | ILLU | 81 | 1 | 1 | 1 | 1 |
| *416516* | SRR11397715 | ILLU | 96 | 1 | 1 | 1 | 1 |
| *430819* | SRR11667145 | ILLU | 13 | 1 | 0.999 | 1 | 0.75 |
| *430820* | SRR11667146 | ILLU | 25 | 1 | 0.999 | 1 | 0.714 |

MALVIRUS scored a perfect precision (100%) on both reference and alternate alleles, while recall on the reference is almost perfect (99.9-100%) with some loss of recall on the alternate alleles. This loss of recall on the alternate alleles is caused by the fact that, especially on ONT data, some SNPs exhibit an unexpected and extremely low coverage that together with the high error rate makes them harder to correctly genotype. A careful inspection of these cases showed that a different choice of parameters (especially the *k*-mer size) improves its accuracy, allowing it to correctly genotype most of these low-covered SNPs at the cost of slightly lower precision. However, we believe that the default parameters of MALVIRUS allow to achieve the best trade-off between precision and recall. Finally, a single SNP (5508:T>C) is unique to the specific strain considered (GISAID ID *EPI_ISL_416410)* and cannot be present in the variant catalog built by the first module of MALVIRUS. Therefore, that variant could not be genotyped by the second module of MALVIRUS. However, since the rapidly increasing number of available complete sequences will broaden the variant catalog, we can expect that this situation will be uncommon in the next few months. On the other hand, such an increasing amount of data does not significantly challenge MALVIRUS since each step of the pipeline is efficient.

We ran MALVIRUS using 8 threads and the analysis of each sample completed in 50/60 minutes requiring less than 7GB of RAM. Such amount of resources is nowadays available on any computer, allowing MALVIRUS to run even on laptops and desktop machines. The first module of our application (catalog creation) required less than 15 minutes and less than 12GB of RAM. Anyway, we point out that the catalog creation needs to be run only when new strains are available, that each catalog can be reused multiple times, and that the software is distributed with a precomputed variant catalog built using the sequences available on NCBI.

## Conclusions

In this work, we presented MALVIRUS, a web application for quickly genotyping viral strains. As shown by our tests, MALVIRUS is able to efficiently and accurately genotype a newly sequenced SARS-CoV-2 strain both from short (Illumina) and long (Oxford Nanopore) reads. Since MALVIRUS benefits from comprehensive variant catalogs, the constantly increasing number of available strains will broaden the completeness of the current variant knowledge, thus boosting the overall accuracy of our pipeline.

## Supporting information

GISAID Acknowledgement Table

## Acknowledgements

This project has received funding from the European Union’s Horizon 2020 research and innovation programme under the Marie Skłodowska-Curie grant agreement No. 872539.

