## Supplementary material for "MALVIRUS: an integrated web application for viral variant calling": GISAID Acknowledgement Table

We gratefully acknowledge the following Authors from the Originating laboratories responsible for obtaining the specimens, as well as the Submitting laboratories where the genome data were generated and shared via GISAID, on which this research is based.

All Submitters of data may be contacted directly via www.gisaid.org

| Accession ID | Originating Laboratory | Submitting Laboratory | Authors |
|---|---|---|---|
| EPI_ISL_416410, EPI_ISL_416411, EPI_ISL_416412, EPI_ISL_416413, EPI_ISL_416415 | Victorian Infectious Diseases Reference Laboratory (VIDRL) | Victorian Infectious Diseases Reference Laboratory and Microbiological Diagnostic Unit Public Health Laboratory, Doherty Institute | Caly L., Seemann T., Schultz M., Druce J., Taiaroa, G. |
| EPI_ISL_416514, EPI_ISL_416515, EPI_ISL_416516 | Victorian Infectious Diseases Reference Laboratory (VIDRL) | Victorian Infectious Diseases Reference Laboratory and Microbiological Diagnostic Unit Public Health Laboratory, Doherty Institute | Caly L., Seemann T., Schultz M., Taiaroa, G., Druce J. |
| EPI_ISL_430819 | Center of Scientific Excellence for Influenza Viruses,National Research Centre (NRC), Egypt. | Center of Scientific Excellence for Influenza Viruses,National Research Centre (NRC), Egypt. | Mohamed Ahmed Ali, Ahmed Kandeil, Ahmed Mostafa, Rabeh El-Shesheny, Mahmoud Shehata, Wael Roshdy, Shymaa Showky Ahmed , Amal Naguib, Nancy M. El Guindy, Mokhtar Gomaa, Ahmed El-Taweel, Ahmed E Kayed, Yassmin Moatasim, Omnia Kutkat, Sara Mahmoud, Mina Kamel, Abo Shama, M Noura, Mohamed El Sayes |
| EPI_ISL_430820 | Center of Scientific Excellence for Influenza Viruses, National Research Centre (NRC), Egypt. | Center of Scientific Excellence for Influenza Viruses, National Research Centre (NRC), Egypt. | Mohamed Ahmed Ali, Ahmed Kandeil, Ahmed Mostafa, Rabeh El-Shesheny, Mahmoud Shehata, Wael Roshdy, Shymaa Showky Ahmed , Amal Naguib, Mokhtar Gomaa, Ahmed El-Taweel, Ahmed E Kayed, Yassmin Moatasim, Omnia Kutkat, Sara Mahmoud, Mina Kamel, Abo Shama, M Noura, Mohamed El Sayes, Nancy M. El Guindy |
